# The chromosome-scale genome assembly of the yellowtail clownfish *Amphiprion clarkii* provides insights into melanic pigmentation of anemonefish

**DOI:** 10.1101/2022.07.21.500941

**Authors:** Billy Moore, Marcela Herrera, Emma Gairin, Chengze Li, Saori Miura, Jeffrey Jolly, Manon Mercader, Michael Izumiyama, Erina Kawai, Timothy Ravasi, Vincent Laudet, Taewoo Ryu

## Abstract

Anemonefish are an emerging group of model organisms for studying genetic, ecological, evolutionary, and developmental traits of coral reef fish. The yellowtail clownfish *Amphiprion clarkii* possesses species-specific characteristics such as inter-species co-habitation, high intra-species color variation, no anemone specificity, and a broad geographic distribution, that can increase our understanding of anemonefish evolutionary history, behavioral strategies, fish-anemone symbiosis, and color pattern evolution. Despite its position as an emerging model species, the genome of *A. clarkii* is yet to be published. Using PacBio long-read sequencing and Hi-C chromatin capture technology, we generated a high-quality chromosome-scale genome assembly initially comprised of 1,840 contigs with an N50 of 1,203,211 bp. These contigs were successfully anchored into 24 chromosomes of 843,582,782 bp and annotated with 25,050 protein-coding genes encompassing 97.0 % of conserved actinopterygian genes, making the quality and completeness of this genome the highest amongst all published anemonefish genomes to date. Transcriptomic analysis identified tissue-specific gene expression patterns, with the brain and optic lobe having the largest number of expressed genes. Further analyses revealed higher copy numbers of *erbb3b* (a gene involved in melanophore development) in *A. clarkii* compared to other anemonefish, thus suggesting a possible link between *erbb3b* and the natural melanism polymorphism observed in *A. clarkii*. The publication of this high-quality genome, along with *A. clarkii*’s many unique traits, position this species as an ideal model organism for addressing scientific questions across a range of disciplines.

## INTRODUCTION

Anemonefish are a group of 28 species that belong to the Pomacentridae family (Fautin et al., 1992). They are social fish that change sex (from male to female) and live in association with sea anemones (Fautin, 1991; Fautin & Allen, 1997). Their visual appeal and ability to breed in captivity initially made them very popular within the ornamental fish trade (Militz et al., 2018; Rhyne et al., 2017). However recently, anemonefish have gained interest from the scientific community as an emerging model species (Roux et al., 2020), providing an alternative to freshwater teleost models such as zebrafish, or larger aquaculture based marine models such as salmon or tuna. This interest has arisen as anemonefish have multiple characteristics that make them attractive future model species for exploring scientific questions across ecological, evolutionary, and developmental fields (reviewed in Roux et al., 2020; Laudet & Ravasi, 2022).

Much of the previous research on anemonefish has been conducted on a limited number of species, namely *Amphiprion ocellaris* and *Amphiprion percula* (Mitchell et al., 2021; Salis et al., 2021). However, the yellowtail clownfish *Amphiprion clarkii* (Bennett, 1830) has unique features that arguably make it the most interesting model species within the Amphiprioninae subfamily (Figure 1a, b). These features include: (1) *A. clarkii* are commonly observed inhabiting anemones with other species of anemonefish (Camp et al., 2016; De Brauwer et al., 2016; Hattori, 2002). While such a trait is not unique to *A. clarkii* (De Brauwer et al., 2016), anemonefish cohabitation primarily involves *A. clarkii* and another species of anemonefish (e.g., *Amphiprion perideraion* or *Amphiprion sandaracinos)* (Camp et al., 2016; Hattori, 2002; Hayashi et al., 2018). Studies of this co-existence can provide further insights into the social structure, complex behaviours, and competitive interactions of anemonefish. (2) As the least host-specific anemonefish, and only inhabitant of *Cryptodendrum adhaesivum* and *Heteractis malu* (Fautin & Allen, 1997; Fautin et al., 1992), *A. clarkii* can provide unique insights into the behavioural and biochemical links between host anemones and resident anemonefish (Da Silva & Nedosyko, 2016). (3) The broad distribution of *A. clarkii*, from Australia to Japan to the Persian Gulf (Fautin et al., 1992), and its wide temperature tolerance (Moyer, 1980) make it a robust and accessible study organism.

**Figure 1.**
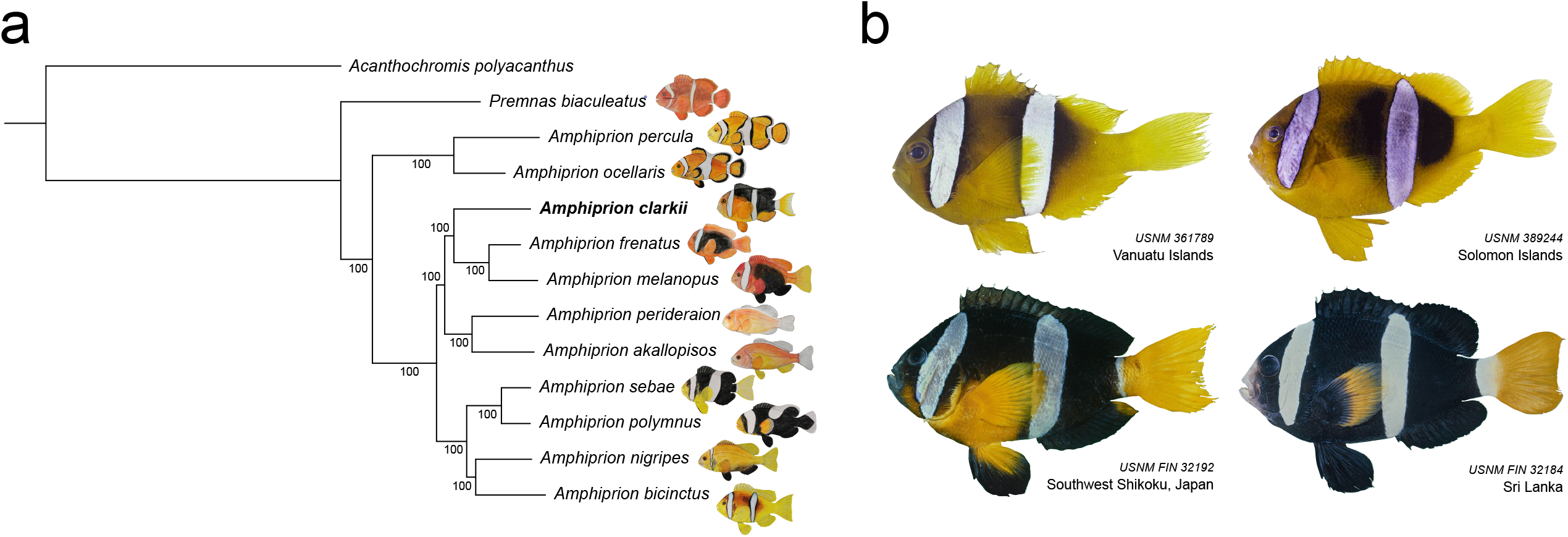
**(a)** Phylogenetic reconstruction of the Amprihprioninae species tree using a maximum-likelihood approach. Bootstrap support values (%) are shown in each branching node. **(b)** Melanistic polymorphism in *Amphiprion clarkii*. Images taken from the Division of Fishes Collections of the Smithsonian National Museum of National History (https://collections.nmnh.si.edu/search/fishes/). Catalog number and sampling location are indicated for each specimen.

In addition to these attributes, *A. clarkii* displays the greatest intra-species variation in colour pigementation pattern of all anemonefishes. Particularly the occurrence of melanism, a darkening (blakish appearance) of body pigmentation that can extend to the anal, pelvic, and dorsal fins (Figure 1b) (Militz et al., 2016). Like other polymorphisms in coral reef fish, melanin-based coloration in *A. clarkii* has been observed to vary with social rank, with increased melanism displaying dominance and sexual dichromatism (Moyer, 1976, 1980) as well as with local variations in habitat (e.g., temperature) (Bell et al., 1982), and host anemone (Fautin & Allen, 1997; Militz et al., 2016). Several species of anemonefish, including *A. clarkii*, exhibit polymorphic melanistic morphs when present in *Stichdactyla* versus *Heteractis* anemones (Militz et al., 2016). Indeed, a recent study (Salis et al., 2021) showed that the developmental timing of white bar formation in juvenile clownfish depends on the anemone species to which they have recruited. Furthermore, it has also been hypothesized that melanism in anemonefish may serve a protective function against the toxicity of host anemones (Maytin et al., 2018). Melanin is an integral component of insect immune responses (Sugumaran, 2002), and there is ample evidence linking melanin pigments and immunity (i.e., phagocytosis, lysosomal enzyme activity, defence against oxidative stress) in vertebrates (Mackintosh, 2001; McGraw, 2005). Thus, it is noteworthy that *A. clarkii* shows a high degree of melanism polymorphism (Figure 1b) and is the only anemonefish found in association with all ten species of sea anemone (with varying levels of toxicity). Altogether, varying levels of melanism within natural populations of anemonefish seems to be the result of environmental cues, host species, and social interactions. However, the extent to which these variables influence this phenomenon is still not fully understood, especially at the molecular level. Based on it’s unique pigmentation pattern, as well as the many other species-specific and genus-specific traits, we anticipate *A. clarkii* will be the focus of future research encompassing a range of scientific disciplines.

For all model species a high-quality genome is an essential resource, required for many advanced genomic approaches. To date, genomes of varying quality have been published for more than ten species of anemonefish, with species such as *A. ocellaris* having multiple genomes available (Lehmann et al., 2019; Marcionetti et al., 2018, 2019; Ryu et al., 2022; Tan et al., 2018). Previous molecular studies of anemonefish have provided novel insights into anemonefish hermaphroditism (Casas et al., 2016) and their mutualistic relationship with anemones (Marcionetti et al., 2019), highlighting the value of genomic data in studies of anemonefish. Yet, the genome of *A. clarkii* is yet to be published, with only a mitochondrial genome currently available (Tao et al., 2016; Thongtam Na Ayudhaya et al., 2019). Moreover, previous genomic studies of *A. clarkii* such as those using ddRADseq (Catalano et al., 2021) and RNA-seq have utilized suboptimal *de novo* assembly-based approaches during analysis. Thus, the availability of a high-quality genome will enhance the appeal and quality of future genetic studies of *A. clarkii*.

Here, we present the first chromosome-scale genome assembly for the yellowtail clownfish *A. clarkii* from Okinawa, Japan. Using a combination of PacBio and Hi-C sequencing, we generated a *de novo* assembly consisting of 1,840 contigs with an N50 of 1,203,211 bp that were successfully anchored into 24 chromosomes of 843,582,782 bp. We annotated 25,050 protein-coding genes encompassing 97.0 % of conserved actinopterygian genes, making the quality and completeness of this *A. clarkii* genome the best of all published anemonefish genomes to date (Lehmann et al., 2019; Marcionetti et al., 2018, 2019; Ryu et al., 2022; Tan et al., 2018) and in the upper echelons of all previously published fish genomes (Conte et al., 2017; Kang et al., 2020; Li et al., 2022; Zhou et al., 2019). Using a comparative genomic approach we also studied genes involved in pigmentation and identified higher copy numbers of the *erbb3b* gene in *A. clarkii*. Interestingly, *erbb3b* encodes for an epidermal growth factor receptor (EGFR)-like tyrosinase kinase (Stein & Staros, 2006) that has been shown to play a crucial role in the establishment of melanocytes (Hultman et al., 2009), thus suggesting a possible link between this gene and the natural melanism polymorphism in *A. clarkii*. Ultimately, the publication of this genome provides a fundamental high-quality resource that will enhance the use of *A. clarkii* as a model species, thus facilitating scientific research that spans a wide range of biological disciplines.

## MATERIALS AND METHODS

### Fish collection and nucleic acid sequencing

Two adult *A. clarkii* (1 male and 1 female) anemonefish were collected for genome and transcriptome sequencing from Tancha Bay, Okinawa (26.4736 N, 127.8278 E) on the 18^th^ of August 2020. These two fish resided together in the anemone *Heteractis crispa* at a depth of 7 m. Following collection, the fish were transferred to Okinawa Institiute of Science and Technology (OIST) Marine Science Station where they remained under natural conditions in a 270 L flow through outdoor tank overnight until they were euthanized the day after. Additionally, ten *A. clarkii* juveniles were collected for quantitative real-time PCR (qPCR) assays from various shallow sites around Okinawa (2 to 11 m deep) between August 2021 and June 2022 (Table S1). Five orange juveniles were collected from *Heteractis* sp. host anemones and five black juveniles from *Stichodactyla* sp. The fish were collected using SCUBA and hand nets, before being euthanized in a 200 mg/L Tricaine Methanesulfonate (MS222) solution and preserved in RNAlater. Samples were placed in 4 °C for 48 hrs and then transferred to a -30 °C freezer until RNA extractions were performed. All fish were euthanized following the guidelines outlined by the Animal Resources Section of OIST Graduate University.

For genome sequencing with PacBio (Pacific Biosciences, CA, USA), the liver of the adult female was extracted, snap frozen in liquid nitrogen, and stored at -80 °C. Liver tissue from the adult male, on the other hand, was used for Hi-C sequencing. Thirteen tissues from the same (male) individual were also used for transcriptome sequencing. Finally, total RNA from the whole body of the juveniles was extracted to perform qPCR. Details on extractions and library preparation are provided in Methods S1.

### Chromosome-scale genome assembly

Raw PacBio long reads were assembled *de novo* using Flye v2.9 (Kolmogorov et al., 2019) with the “keep haplotypes” option. Assessment of the resulting genomic contigs with Benchmarking Universal Single-Copy Orthologs (BUSCO) v4.1.4 (Simão et al., 2015) and the Actinopterygii-lineage dataset (actinopterygii_odb10) identified high levels of gene duplication. Therefore, duplicates were removed from the initial Flye assembly using purge_dups v0.03 (Guan et al., 2020). The chromosome-scale genome assembly was generated by Phase Genomics using the *de novo* assembly, FALCON-phase (Kronenberg et al., 2018), Hi-C sequencing reads, and Phase Genomics’ Proximo algorithm based on Hi-C chromatin contact maps (as described in Bickhart et al., 2017). Error-correction of this chromosome-scale assembly was conducted with Illumina short reads and Pilon v1.23 (Walker et al., 2014). Quality-trimmed Illumina short reads (Trimmomatic v0.39) (Bolger et al., 2014) using the parameters “ILLUMINACLIP:TruSeq3-PE.fa:2:30:10:8:keepBothReads LEADING:3 TRAILING:3 MINLEN:36” were aligned to the genome using Bowtie2 v2.4.1 (Langmead & Salzberg, 2012) with the default parameters, and the resulting SAM files were converted to BAM format using SAMtools v1.10 (Li et al., 2009). BAM files were then used as input for error correction with Pilon. The quality and completeness of the final assembly was assessed using Quast v5.0.2 (Mikheenko et al., 2018) and BUSCO v4.1.4 (actinopterygii_odb10) (Simão et al., 2015).

### Genome size and coverage estimation

Genome size and heterozygosity were estimated using quality-trimmed Illumina short reads (as described above), Jellyfish v2.3.0 (Marçais & Kingsford, 2011) with k-mer = 17, and GenomeScope v1.0 (Vurture et al., 2017) with default parameters. Additionally, overall mean genome-wide base level coverage of the final assembly was calculated by aligning the raw PacBio reads to the assembled chromosome sequences using Pbmm2 v1.4.0 (https://github.com/PacificBiosciences/pbmm2). The genomeCoverageBed function of BEDTools v2.30.0 (Quinlan, 2014) was then used to calculate the per-base coverage of aligned reads across all chromosomal sequences.

### Prediction of gene models in *A. clarkii*

Repetitive elements were identified *de novo* using RepeatModeler v2.0.1 (Flynn et al., 2020) with the “LTRStruct” option. RepeatMasker v4.1.1 (Tempel, 2012) was used to screen known repetitive elements with two inputs: (1) the RepeatModeler output and (2) the vertebrata library of Dfam v3.3 (Storer et al., 2021). The resulting output files were validated and merged before redundancy was removed using GenomeTools v1.6.1 (Gremme et al., 2013). To identify and annotate candidate gene models, BRAKER v2.1.6 (Brůna et al., 2021) was used with mRNA and protein evidence. For annotation with BRAKER, the chromosome sequences were soft masked using the maskfasta function of BEDTools v2.30.0 (Quinlan, 2014) with the “soft” option. Protein evidence consisted of protein records from UniProtKB/Swiss-Prot (UniProt Consortium, 2021) as of January 11, 2021 (563,972 sequences) as well as selected fish proteomes from the NCBI database (*A. ocellaris*: 48,668, *Danio rerio*: 88,631, *Acanthochromis polyacanthus*: 36,648, *Oreochromis niloticus*: 63,760, *Oryzias latipes*: 47,623, *Poecilia reticulata*: 45,692, *Stegastes partitus*: 31,760, *Takifugu rubripes*: 49,529, and *Salmo salar*: 112,302). Transcriptomic reads from 13 tissues were used as mRNA evidence. These Illumina short reads were trimmed with Trimmomatic v0.39 (Bolger et al., 2014) as described above and mapped to the chromosome sequences with HISAT2 v2.2.1 (Kim et al., 2019). The resulting SAM files were converted to BAM format with SAMtools v1.10 (Li et al., 2009) and used as input for BRAKER. Of the resulting gene models, only those with supporting evidence (mRNA or protein hints) or with homology to the Swiss-Prot protein database (UniProt Consortium, 2021) or Pfam domains (Mistry et al., 2021) were selected as final gene models. Homology to Swiss-Prot protein database and Pfam domains was identified using Diamond v2.0.9 (Buchfink et al., 2015) or InterProScan v5.48.83.0 (Zdobnov and Apweiler, 2001), respectively. Functional annotation of the final gene models was completed using NCBI BLAST v2.10.0 (Altschul et al., 1990) with the NCBI non-redundant (nr) protein database. Gene Ontology (GO) terms were assigned to *A. clarkii* genes using the BLAST output and the “gene2go” and “gene2accession” files from the NCBI ftp site (https://ftp.ncbi.nlm.nih.gov/gene/DATA/). Completeness of the gene annotation was assessed with BUSCO v4.1.4 (actinopterygii_odb10) (Simão et al., 2015).

### Mitochondrial genome assembly and annotation

Quality trimmed Illumina reads were used as input for GetOrganelle v1.7.0 (Jin et al., 2020) which was used to assemble the mitochondrial genome of *A. clarkii*. Mitochondrial genes were then annotated with MitoAnnotator v3.67 (Sato et al., 2018). The mitochondrial genome assembled here was compared to two previously published mitochondrial genomes of *A. clarkii* (Tao et al., 2016; Thongtam Na Ayudhaya et al., 2019) (NCBI accessions: NC_023967.1 and AB979449.1) using BLASTn v2.10.0 (Altschul et al., 1990) with an e-value 10^−4^ as a threshold to predict overall sequence identity.

### Analysis of tissue-specific gene expression

As for gene annotation, quality trimmed transcriptomic reads from 13 tissues were mapped to the chromosome assembly with HISAT2 v2.2.1 (Kim et al., 2019) and the resulting SAM files were converted to BAM format using SAMtools v1.10 (Li et al., 2009). The resulting BAM files and final gene annotation file were used as input into StringTie v2.1.4 (Pertea et al., 2016) to quantify expression levels and normalize TPM (transcripts per million). The tissue specificity index (*τ*) of each gene was calculated using the R package tispec v0.99 (Condon, 2020) and a two-dimensional histogram was used to display the relationship between *τ* and expression level (TPM). The number of genes expressed in each tissue and different combinations of tissues were displayed in an Upset plot generated with the UpSetR v1.4.0 R package (Conway et al., 2017).

### Gene orthology and phylogenetic analyses

Orthologous relationships between *A. clarkii* and the other anemonefish were investigated using OrthoFinder v2.5.2 (Emms & Kelly, 2019). Briefly, protein sequences of *A. clarkii, Amphiprion akallopisos, Amphiprion bicinctus, Amphiprion frenatus, Amphiprion melanopus, Amphiprion nigripes, Amphiprion ocellaris, Amphiprion percula, Amphiprion perideraion, Amphiprion polymnus, Amphiprion sebae*, and *Premnas biaculeatus* (Lehmann et al., 2019; Marcionetti et al., 2018, 2019; Ryu et al., 2022), and the spiny chromis *A. polyacanthus* (used as the outgroup species), were reciprocally blasted against each other and clusters or othologous genes were defined using the default settings. In all cases, only the longest isoform of each gene model was used. Sequences of single-copy orthologs present in all species were aligned using MAFFT v7.130 (Katoh & Standley, 2013) using the options “local pair”, “maxiterate 1,000”, and “leavegappyregion”, trimmed with trimAl v1.2 (Capella-Gutiérrez et al., 2009) using the “strict” flag, and then concatenated with FASconCAT-G (Kück & Longo, 2014). Maximum-likelihood phylogenetic trees were then constructed with RAxML v8.2.9 (Stamatakis, 2014). The MPI version (raxmlHPC-MPI-AVX) was executed using a LG substitution matrix, heterogeneity model GAMMA, and 1,000 bootstrap inferences. Trees were visualized using iTOL v6.4 (Letunic & Bork, 2021). Branch supports in the trees were evaluated with the standard bootstrap values from RaxML.

### Identification of pigmentation genes

Based on Lorin et al. (2018) and Salis et al. (2021), a list of 208 genes known to be involved in pigmentation were identified for this study (Table S2). For each of these genes, the related protein sequence of *A. ocellaris* (or, if not available, the closest related species) was retrieved from the Ensembl genome database (https://www.ensembl.org, last accessed on February 2022). Next, a BLASTp search (using the parameters “evalue 10^−10^” and “max_target_seqs 5”) of these 208 protein sequences was performed against the *A. clarkii* gene models. These were then confirmed to be the correct pigmentation genes by checking the previously completed *A. clarkii* gene annotation.

### Confirmation of *erbb3b* genes in *A. clarkii*

The presence of three *erbb3b* genes identified in the *A. clarkii* genome was validated using polymerase chain reaction (PCR) and DNA from the same individual used for whole-genome sequencing. Additionally, PCR was used to investigate the presence of *erbb3b* genes in other species of anemonefish (*A. ocellaris, A. frenatus, A. polymnus, A. perideraion, A. sandaracinos*, and *Amphiprion akindynos)*. For these species, DNA was extracted from a piece of caudal fin using a Maxwell® RSC Blood DNA Kit (Promega, Madison, WI, USA). Extractions were performed following the manufacturer’s instructions with the exception of a longer two-hour lysis step. DNA was quantified using a Qubit dsDNA BR (Broad Range) Assay Kit (Thermo Fisher Scientific, Waltham, MA, USA). DNA was then diluted to a working concentration of 20 ng/μL and stored at -30 °C.

Primers targeting a conserved region of intron 8 of all three *erbb3b* genes in *A. clarkii* were designed using Geneious v2022.1 (Biomatters). Gaps of different lengths were present across the three genes (Figure S1), thus making it easy to amplify them using only one pair of primers. PCRs were performed using the forward 5’ TGTCCACTTCCAGGATGAGAC 3’ and reverse 3’ ACCCCTCGATCTCATCTCTGT 5’ primers. Each PCR was run using 12.5μL Q5® High-Fidelity 2X Master Mix (New England Biolabs, Ipswich, MA, USA), 2.5μL template DNA, 1.25μL 10μM forward and reverse primer, and 7.5μL nuclease-free water for a final reaction volume of 25 μL. The thermal cycling conditions used were 30 s 98 °C, followed by 35 cycles of 10 s at 98 °C, 30 s at 67 °C, and 30 s at 72 °C, followed by a final extension step of 2 min at 72 °C. For each sample, PCR products were visualized using 2 % agarose gel electrophoresis (Figure S2), excised from the gel and purified using a QIAquick Gel Extraction Kit (Qiagen, Hilden, Germany). PCR amplicons were then bidirectionally sequenced by the company FASMAC, which uses Applied Biosystems Big Dye Terminator v3.1 technology and an Applied Biosystems 3130xl Genetic Analyzer (Applied Biosystems, Waltham, MA, USA). Sequence analysis was performed using the software Geneious v2022.1 (Biomatters).

### qPCR assays to measure *erbb3b* gene expression in *A. clarkii*

In total, ten juveniles (five orange and five black) were assayed to measure gene expression of the three *erbb3b* genes previously identified in *A. clarkii*. Specific primers for each *A. clarkii erbb3b* gene (two short genes containing 1,911 bp and one long gene of 4,275 bp) were designed manually based on their genomic sequence (Table S3). Primers previously used by Roux et al. (2022) with *A. ocellaris* were used to target the housekeeping genes ribosomal protein L7 (*rpl7*) and ribosomal protein L32 (*rpl32*). Extracted RNA from each juvenile (as described in Methods S1) was converted to cDNA using PrimeScript RT-PCR Kit (Takara Bio, Shiga, Japan). The efficiency and specificity of the designed primers was tested through PCR using the GoTaq Green Master kit (Promega, Madison, USA) with thermal cycling conditions of 2 min at 95 °C, followed by 30 cycles of 45 s at 95 °C, 45 s at 60/63/65 °C, and 30 s 72 °C, a final extension step of 5 min at 72 °C, preservation at 4 °C, and subsequent agarose gel electrophoresis (Figure S3). The specificity was also tested through direct forward and reverse Sanger sequencing by aligning the forward and reverse outputs and blasting the obtained amplicons against the reference genomic sequences (Figure S4).

The expression of each *erbb3b* gene and the two housekeeping genes (*rpl7* and *rpl32*) was obtained by RT-qPCR at 65 °C (PrimeScript transcriptase, Takara, SYBRgreen) and normalized with the Pfaffle equation (Ståhlberg et al., 2004):

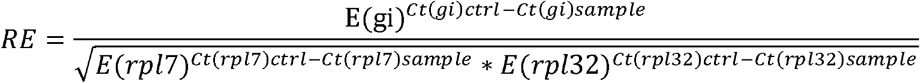

where RE is the relative expression, E(x) is the efficiency of the amplication for gene x, and Ct(x) is the quantification cycle of gene x.

## RESULTS AND DISCUSSION

### Chromosome-scale genome assembly of *A. clarkii*

We assembled the genome of the anemonefish *A. clarkii* by sampling two individuals from Okinawa and generating 19,675,845 PacBio reads with an average read length of 13,144 bp (Table S4). These reads were assembled *de novo* using Flye v2.9 (Kolmogorov et al., 2019) with the initial assembly consisting of 2,635 contigs of 855,782,104 bp with an N50 of 1,187,902 bp. Following processing with Purge_Dups v0.0.3 (Guan et al., 2020) the final *de novo* assembly consisted of 1,840 contigs of 845,361,362 bp and had an N50 of 1,203,211 bp. Using 228,099,434 150 bp Hi-C reads from liver tissue and the ProximoTM scaffolding platform (Phase Genomics, WA, USA), we generated 24 chromosomes of 843,295,090 bp and 168 short scaffolds (2,826,673 bp) that were not placed into chromosomes. This chromosome-scale assembly was polished with Illumina short reads using Pilon (Walker et al., 2014) generating a final assembly of 843,582,782 bp. Chromosome lengths ranged from 42,519,526 bp to 20,115,265 bp (Figure 2). The mean base-level coverage of these chromosomes was 250.4 x. The final *A. clarkii* genome contained 118,106 non-ATGC characters, a GC content of 39.71 % and a repeat content of 44.26 % (Table 1). The structure of our genomic assembly was compared to properties of the *A. clarkii* genome estimated by Jellyfish v2.3.0 (Marçais & Kingsford, 2011) and GenomeScope v1.0 (Vurture et al., 2017) with Illumina short reads. At kmer = 17, genome size was estimated at 793,832,155 bp, repeat content was estimated at 42.33 % and heterozygosity was estimated at 0.51 %. Repeat content was identified using RepeatMasker v4.1.1 (Tempel, 2012) by querying repetitive elements from the Dfam (Storer et al., 2021) vertebrata library and repetitive elements identified *de novo* using RepeatModeler v2.0.1 (Flynn et al., 2020) against the *A. clarkii* genome. This approach identified repeat content of 373,358,331 bp (Figure S5). Of the identified repetitive elements, DNA transposons were the most frequent, occupying 23.26 % of the *A*.*clarkii* genome. Long interspersed nuclear elements (7.33 %), long terminal repeats (3.75 %), and simple repeats (1.73 %) were the next most frequent in the genome. However, 27.93 % of the *A*.*clarkii* genome is occupied by repetitive elements that could not be identified (Figure S5).

**Figure 2.**
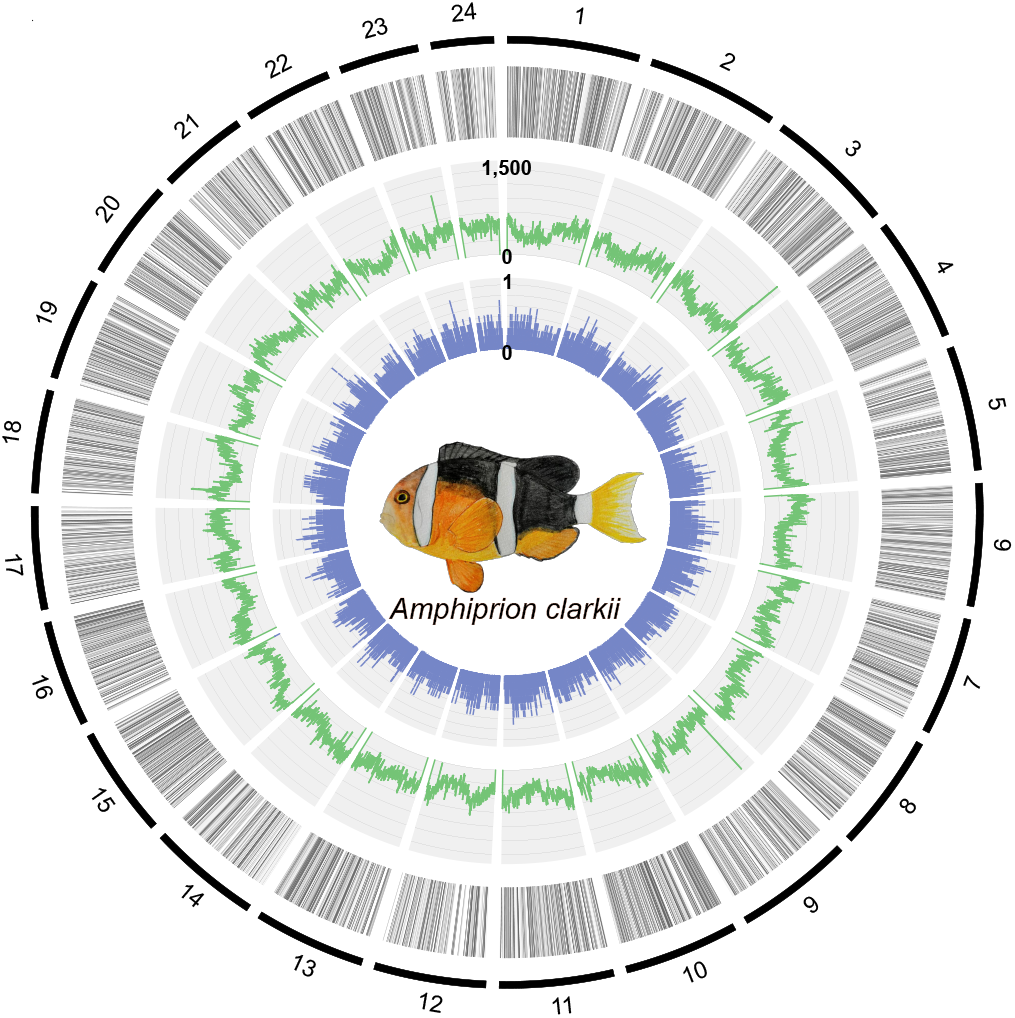
The genome structure of *Amphiprion clarkii*. From the outside in, the circos plot layers display: (1) chromosome order and size, (2) genic regions of the genome, (3) number of repeats per 100 kb, and (4) tissue specificity index of genes displayed in layer 1. Drawing of *A. clarkii* is displayed in the middle of the circos plot.

**Table 1.** Genome assembly statistics, gene annotation statistics, and BUSCO completeness.

|  |  |
| --- | --- |
| **Chromosome assembly size** | **843,582,782 bp** |
| Non-ATGC characters | 118,106 |
| GC content | 39.7 % |
| Mean base-level coverage | 250.4 x |
| Repeat content | 44.3 % |
| Chromosome-scale N50 | 26,694,648 bp |
| Contig N50 | 1,203,211 bp |
| **BUSCO genome completeness** | **3,593 (98.7 %)** |
| Complete and single copy | 3,560 (97.8 %) |
| Complete and duplicated | 33 (0.9 %) |
| Fragmented | 13 (0.4 %) |
| Missing | 34 (0.9 %) |
| Number of protein coding genes | 25,050 |
| **BUSCO gene annotation completeness** | **3,532 (97.0 %)** |
| Complete and single copy | 3,498 (96.1 %) |
| Complete and duplicated | 34 (0.9 %) |
| Fragmented | 41 (1.1 %) |
| Missing | 67 (1.9 %) |

Comparison with the two other chromosome-scale anemonefish genomes revealed similar structures and assembly statistics between the *A. clarkii, A. percula*, and *A. ocellaris* genomes (Lehmann et al., 2019; Ryu et al., 2022). For example, with sizes of 890,200,000 bp and 856,612,077 bp, respectively, the *A. percula* and *A. ocellaris* genomes are only slightly larger than the 843,582,782 bp *A. clarkii* genome assembled here, whilst the GC content of all three genomes is between 39.55 and 39.71 %. The repeat content of the *A. clarkii* genome closely matched that of *A. ocell*aris (44.7 % vs 44.26 %), yet it was greater than that of the *A. percula* genome (28 %). This is likely due to the different repeat annotation methods used by Lehmann et al. (2018) compared to those used here and by Ryu et al. (2022). Similarities in repeat content of the sister species *A. ocellaris* and *A. percula* (Litsios et al., 2014) would be expected to match the similarity between *A. ocellaris* and *A. clarkii*. Furthermore, the characteristics of these high-quality anemonefish genomes match that of closely related species within the Pomacentridae family such as *A. polyacanthus* (991,600,000 bp) (ASM210954v1, GCF_002109545.1, NCBI).

Genome completeness was assessed using BUSCO v4.1.4 (Simão et al., 2015) and the Actinopterygii-lineage dataset. The *A. clarkii* genome contained 3,593 conserved actinopterygian benchmark genes giving a BUSCO score of 98.7 % (Complete and single copy: 97.8 %; Complete and duplicated: 0.9 %; Fragmented: 0.4 %; Missing: 0.9 %) (Table 1). Previous non chromosome-scale anemonefish genomes (Marcionetti et al., 2018, 2019; Tan et al., 2018) are much less contiguous (contig numbers of 17,801 and 6,404, respectively) and have a maximum BUSCO score of 96.5 % (Actinopterygii). Although contiguity is important, the genic completeness of an assembly is vital for its future use by the research community. With BUSCO scores of < 97.1 % previous chromosome-scale anemonefish genome assemblies are less complete than the assembly presented here, highlighting this *A. clarkii* assembly as the best quality for anemonefish to date.

### *A. clarkii* gene annotation

The genome was annotated using BRAKER v2.1.6 (Brůna et al., 2021) with mRNA and protein evidence. This resulted in an initial 41,083 predicted gene models. These gene models included different isoforms from the same gene locus, therefore gene models were filtered to keep only the longest isoform of each gene. This resulted in 36,949 unique gene models. Only gene models with either mRNA or protein evidence support (24,571) or homology to the Swiss-Prot protein database or Pfam domains (479) were retained. This resulted in 25,050 final gene models. Of these 25,050 gene models 23,700 (94.61 %) had significant homology to the NCBI *nr* database (bit-score ≥ 50) and 19,982 genes (79.77 %) had at least one associated GO term. The completeness of this set of annotated genes was assessed using BUSCO v4.1.4 (Simão et al., 2015) and the Actinopterygii-lineage dataset. The 25,050 gene models contained 3,532 conserved actinopterygian benchmark genes, giving a BUSCO score of 97.0 % (Complete and single copy: 96.1 %; Complete and duplicated: 0.9 %; Fragmented: 1.1 %; Missing: 1.9 %) (Table 1). Of the previously reported anemonefish genome annotations, the annotation of the chromosome-scale *A. ocellaris* genome (Ryu et al., 2022) was the most complete with a BUSCO score of 96.62 %. Thus, the annotation reported here represents the most complete genome annotation for an anemonefish to date. This high-quality annotation will facilitate genetic studies of *A. clarkii* that require an understanding of specific gene functions and locations.

### Assembly and annotation of mitochondrial genome

The mitochondrial genome of *A. clarkii* was assembled using GetOrganelle v1.7.0 (Jin et al., 2020) and annotated with MitoAnnotator v3.67 (Sato et al., 2018). This resulted in a 16,812 bp circular mitogenome that contained 37 organelle genes consisting of 13 protein-coding genes, 22 tRNAs, and two rRNAs, as well as one control region (Figure S6, Discussion S1).

### Gene expression tissue specificity

The tissue specificity of the 25,050 *A. clarkii* genes identified here was investigated using the transcriptomes of 13 different tissues (Table S4). Total number of genes and unique number of genes expressed per tissue, as well as tau index (*τ*) (Kryuchkova-Mostacci & Robinson-Rechavi, 2017) were used to quantify tissue specificity. A total of 1,814 genes were expressed in all tissues (Figure 3a), which is similar to the 1,957 genes expressed in all tissues in *A. ocellaris* (Ryu et al., 2022), but less than the ∼8,000 genes ubiquitously expressed in multiple human and mouse tissues (Ramsköld et al., 2009).

**Figure 3.**
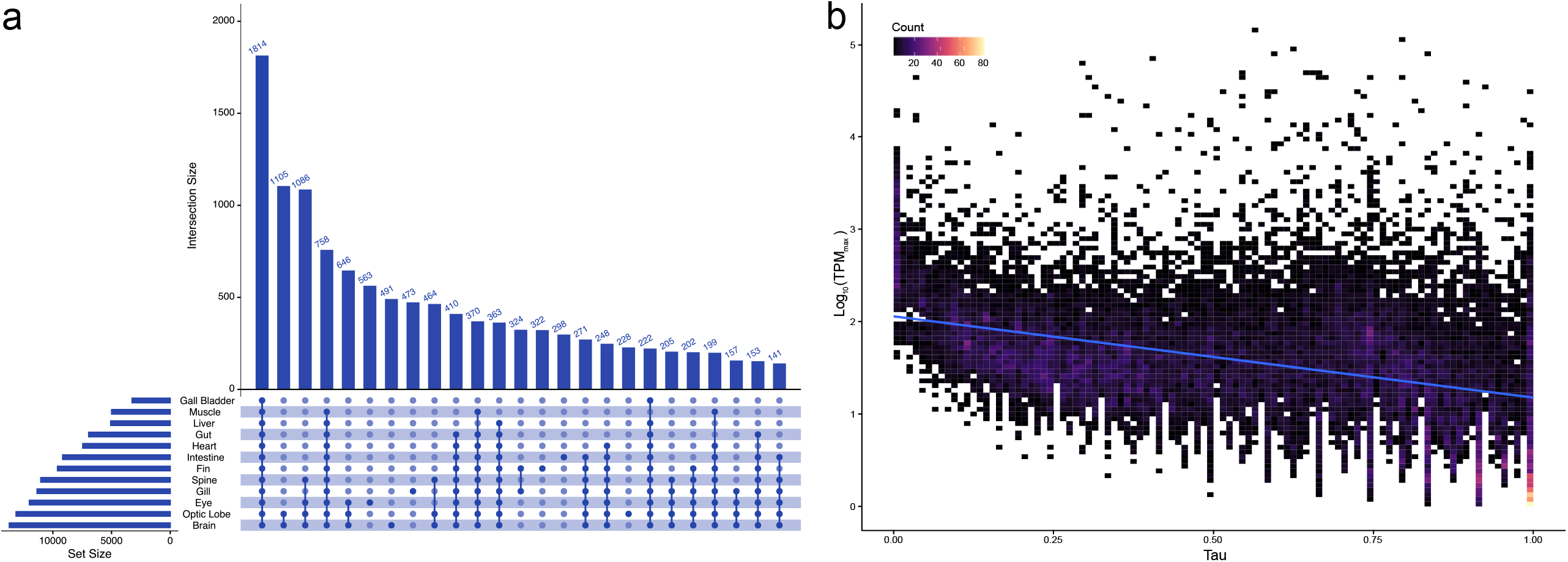
**(a)** Upset plot displaying the number of genes expressed (intersection size) in individual and combinations of different tissues. Transcripts per million (TPM) values of > 10 were used as a threshold for gene expression. Note that “rest of brain” and “cerebrum” tissues are combined to give the final “brain” tissue counts. **(b)** Two-dimensional histogram displaying the relationship between the maximum TPM and tissue specificity index (Tau, *τ*) of each gene. Trendline displays Pearson’s correlation between *τ* and log_10_.

Although only 1,814 genes were expressed in all tissues, 3,697 genes have a *τ ≤* 0.2, indicating they are expressed in nearly all tissues without biased expression, and are therefore considered housekeeping genes. Thus, the number of housekeeping genes identified in *A. clarkii* is very similar to *A. ocellaris* (3,431 housekeeping genes) (Ryu et al., 2022). Genes with greater tissue specificity were more abundant than housekeeping genes as we identified 4,362 highly specific genes (0.85 *≤ τ* < 1) as well as 1,068 absolutely tissue specific genes (*τ* = 1), that were only expressed in one tissue type. The eye expressed the highest number of these unique genes (563) with the brain (491) displaying the second highest (Figure 3a).

However, when considered together, the brain and optic lobe expressed 1824 absolutely tissue specific genes. The number of unique genes expressed in tissue types reflected the total number of genes expressed in different tissues, as the brain (13,714), optic lobe (13,138), and eye (12,003) expressed a high number of genes. The number of genes expressed per tissue are very similar to those observed for *A. ocellaris* (Ryu et al., 2022) and the corresponding human (Ramsköld et al., 2009) tissues, yet is slightly higher than the number expressed in corresponding rainbow trout tissues (Salem et al., 2015). Interestingly, the gall bladder expressed the lowest number of genes (3,252), with only 19 of these being absolutely tissue specific. Across all tissues, tissue specificity of gene expression negatively correlated (Pearson’s correlation coefficient between *τ* and log_10_) with expression levels (Figure 3b), indicating that tissue-specific genes have lower expression levels in general (Kryuchkova-Mostacci & Robinson-Rechavi, 2017).

### Ortholog identification and Anemonefish phylogeny

We used OrthoFinder v2.5.2 (Emms & Kelly, 2019) to identify orthologous relationships between the amino acid sequences of *A. clarkii* and 11 other anemonefish (and the spiny chromis *A. polyacanthus* as an outgroup species). Overall, 96.7% of the sequences could be assigned to one of 29,855 orthogroups, with the remainder identified as “unassigned genes” with no clear orthologs (Table S5). Fifty percent of all proteins were in orthogroups consisting of ≥ 13 genes and were contained in the largest 10,641 orthogroups. 15,771 orthogroups were shared amongst all the species examined here, of which 12,600 consisted entirely of single-copy genes (Table S5).

Phylogenetic reconstruction using these single-copy genes yielded robust phylogenetic relationships, with all branches supported by 100 % boostrap values (Figure 1a). Furthermore, our tree topology is consistent with previous studies (Litsios et al., 2014; Litsios & Salamin, 2014; Marcionetti et al., 2019). Recovered at the base of the tree was *P. biaculeatus*, with the *A. ocellaris*/*A. percula* complex at the root of all other anemonefish, and four major clades: (1) *A. frenatus* and *A. melanopus* and its sister species *A. clarkii*, (2) the skunk anemonefishes *A. akallopisos* and *A. perideraion*, (3) the closely related species *A. polymnus* and *A. sebae*, and (4) an Indian Ocean clade represented by *A. bicinctus* and *A. nigripes*. Situated in the inferior half of the tree, *A. clarkii* is neither the most ancestral nor the most derivative species, but a species with an intermediate level of evolution within the Amphiprioninae subfamily (Litsios et al., 2014; Litsios & Salamin, 2014). Noteworthy, this tree differs from the one recently reported in Ryu et al., (2022). Pomacentrids (anemonefishes in particular) have long been a challenge in systematics due to their high diversity and intraspecific variation (Tang et al., 2021), thus future analyses including more species, especially those located close to the base of the tree, might be critically important in establishing a well-resolved phylogeny.

### Identification of specific pigmentation genes in *A. clarkii*

To identify gene families that are enriched in the *A. clarkii* genome, we counted the orthogroups with genes in all species, and then selected those in which numbers were ≥ 2x higher in *A. clarkii* than other anemonefishes. Interestingly, most of these were associated with processes related to melanogenesis (Table S6). Keratin type II (OG0000508) and receptor tyrosine protein kinase *erbb3* (OG0001853) stood out, as these contained 4 and 3 genes in *A. clarkii* compared with 2 and 1 orthologs in other anemonefish species, respectively (however, *A. ocellaris* contained 2 *erbb3* orthologs) (Figure 4a, Table S6). Keratins are major structural proteins in epithelial cells that influence the distribution and arrangement of melanosomes (Gu & Coulombe, 2007), which ultimately impact the color patterning of animals. Indeed, mutations in keratin domains can cause hyper-/hypo-pigmented phenotypes (Uttam et al., 1996). Keratin type II, in particular, has been implicated in the production of color in frogs (in morphs that have black dorsum and legs) (Stuckert et al., 2021). Here, one gene was identified as keratin type II cytoskeletal 8-like isoform, whereas the other three were annotated as keratin type II cytoskeletal cochleal-like (Figure S7), a key component of the large transcellular cytoskeletal network in the cochlea’s organ of Corti (that contributes to hearing) (Mogensen et al., 1998). Interestingly, however, keratin type II cytoskeletal cochleal-like has been found to be highly expressed in trout skin (Djurdjevič et al., 2019). Receptor tyrosine protein kinase *erbb3*, on the other hand, belongs to the epidermal growth factor receptor (EGFR) family of receptor tyrosine kinases (ErbB), a group of proteins that play essential roles in regulating cell proliferation and differentiation (Stein & Staros, 2006; Wieduwilt & Moasser, 2008). In particular, the ErbB gene *erbb3b* signaling is required for the formation of new melanocytes during metamorphosis (Hultman et al., 2009). Indeed, mutations in this gene result in a phenotype lacking most melanophores (i.e., picasso mutant in zebrafish) (Budi et al., 2008).

**Figure 4.**
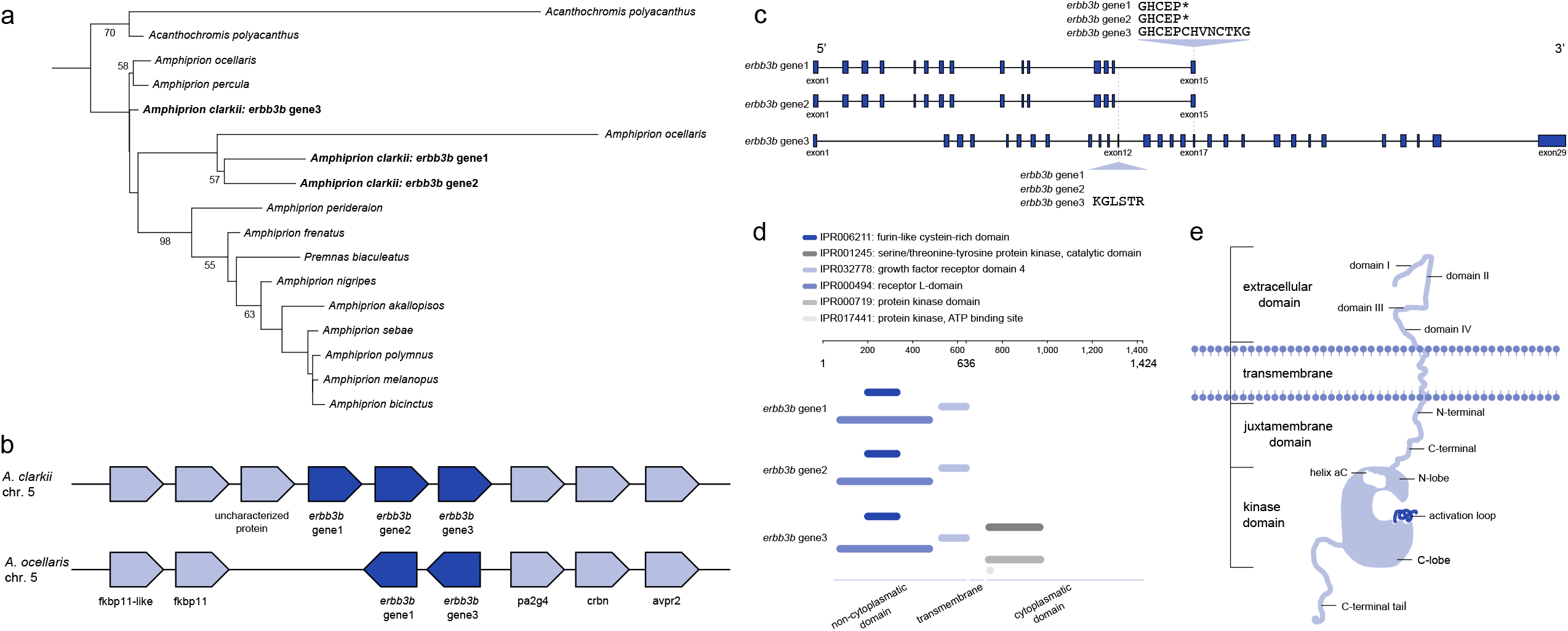
**(a)** Maximum-likelihood phylogeny of protein sequences from the *erbb3b* gene in anemonefish. Boostrap support values (%) above 50 are shown in each branching node. **(b)** Syntenic *erbb3b* genes between *Amphiprion clarkii* and the false clownfish *Amphiprion ocellaris*. The two *erbb3b* genes identified in *A. ocellaris* are orthologous to the *erbb3* gene1 and *erbb3b* gene3 in *A. clarkii*. Dark blue-colored boxes indicate the *erbb3b* gene and light blue-colored boxes represent the flanking orthologous genes. Each gene is arranged considering the transcriptional direction, and the physical distance is ignored. **(c)** Structure of *erbb3b* genes in *A. clarkii*. Both short sequences (*erbb3b* gene1 and *erbb3b* gene2) are comprised of 15 exons whereas the long sequence (*erbb3b* gene3) consists of 29 exons. The short sequences have a gap that corresponds to exon12 of the long sequence and end in the same position as exon17 of the long gene. Exons are represented by blue-colored boxes and introns by a black solid line. Each exon and intron is represented considering the size and physical distance. Asterisk (*) represents the stop codon. **(d)** Functional analysis of the *erbb3b* genes as implemented by InterPro. Protein domains are color-coded in the legend. **(e)** Structure diagram of the erbb3 protein (adapted from Li et al., 2013) including an extracellular ligand binding domain, a transmembrane helix domain, and an intracellular tyrosine kinase domain. The extracellular domain is made of a tandem repeat of leucine-rich (domains I and III) and cysteine-rich segments (domains II and IV). The intracellular domain is a continuation of the transmembrane region and is divided into a juxtamembrane region (which is in turn divided into N-terminal and C-terminal), a kinase domain, and a C-terminal tail. The kinase domain includes the N-lobe, helix aC, activation loop, and C-lobe.

### *erbb3b* genes in *A. clarkii*

Three sequences annotated as *erbb3b* (herein referred to as *erbb3b* gene1, *erbb3b* gene2, and *erbb3b* gene3) were identified in *A. clarkii*, two were identified in *A. ocellaris*, and only one in all the other anemonefish species (Figures 4a and 4b). Notably, *erbb3b* gene1 and *erbb3b* gene2 are much shorter (636 amino acids) compared to *erbb3b* gene3 (1,424 amino acids) (Figure 4c). In the case of *A. ocellaris*, one short and one long sequence was retrieved (Figure 4b). Synteny analysis between *A. clarkii* and *A. ocellaris* revealed all genes are located in tandem on chromosome 5, and are flanked by peptidyl-prolyl cis-trans isomerase FKBP11 and proliferation-associated 2G4 genes (Figure 4b). The protein sequences of *erbb3b* gene1 and *erbb3b* gene2 are almost identical, with both comprising of 15 exons, a gap that corresponds to exon 12 in *erbb3b* gene3, and a stop codon at the same position as exon 17 in the *erbb3b* gene3 (Figure 4c). Furthermore, functional analysis using the InterPro database (Blum et al., 2021) revealed six protein domains: (1) the extracellular growth factor receptor domain IV, (2) the furin-like cysteine-rich domain, (3) the receptor L-domain, and the cytoplasmatic domains (4) serine/threonine-tyrosinase protein kinase, (5) protein kinase (catalytic subunit), and (6) protein kinase ATP binding site (Cho et al., 2002; Hanks et al., 1988). Interestingly, neither *erbb3b* gene1 nor *erbb3b* gene2 have the cytosolic protein kinase domains but only the extracellular ligand domains (Figure 4d).

The structure of *erbb3b* (Figure 4e) is typical in the ErbB receptor tyrosine kinase family. It includes an extracellular ligand binding domain of 600-630 amino acids, a transmembrane helix domain, and an intracellular domain of ∼600 amino acids that includes the tyrosine kinase and regulatory sequences (Li et al., 2013). The extracellular domain itself is made of a tandem repeat of leucine-rich segments that make up the ligand binding (domains I and III), and cysteine-rich domains (II and IV, with the former containing the dimerization arm). The intracellular domain is a continuation of the transmembrane region and is divided into a juxtamembrane region (which is in turn divided into N-terminal and C-terminal), kinase domain, and C-terminal tail. Located in the N-terminal extremity of the catalytic domain, there is a lysine residue that has been shown to be involved in ATP binding. The kinase domain includes an N-lobe, helix αC, activation loop, and C-lobe (Li et al., 2013). However, unlike other ErbB family members, *erbb3* lacks endogenous kinase activity (Jura et al., 2009; Li et al., 2013). Thus, phosphorylation of target proteins only occurs if ligand binding leads to dimerization with other tyrosinase kinase receptors, such as *erbb2*, that do have kinase activity (Jura et al., 2009; Li et al., 2013). Following ligand binding, intracellular pathways are then triggered, in this case, the formation of new melanocytes (Hultman et al., 2009).

Given that *A. clarkii* is a polymorphic species in terms of pigmentation, particularly melanism, finding higher copy numbers of a gene implicated in melanophore development calls for further analysis.

### Validation of *erbb3b* genes in *A. clarkii* and other anemonefish species

Importantly, all three *erbb3b* genes described above were confirmed to be present in the *A*.*clarkii* genome (Figure S2). However, whilst our bioinformatic analysis identified only one ortholog for the other species (except *A. ocellaris*, for which we identified two), PCR and Sanger sequencing highlighted the presence of two *erbb3b* genes in all other anemonefish species tested here. Alignment of these sequences to the *erbb3b* genes from *A. clarkii* indicated that these two copies correspond to one short and one long copy of *erbb3b* (orthologous to *erbb3b* gene1 and *erbb3b* gene3, respectively). Thus, to further validate these results, we performed a BLASTn search (using the parameters “task blastn”, “evalue 10^−10^”, and “max_target_seqs 5”) of the three *A. clarkii erbb3b* genes against the genomes of all the other species (Lehmann et al., 2019; Marcionetti et al., 2018, 2019; Ryu et al., 2022). This resulted in genes matching sequences annotated as receptor tyrosine protein kinase *erbb3* (Table S7). However, these matches corresponded to only 25−35 % of the *A. clarkii erbb3b* genes total length (10,283 bp for *erbb3b* gene1 and 20,883 bp for *erbb3b* gene3 from start to stop codons). With the exception of the *A. ocellaris* and *A. percula* genomes (for which we obtained two and one match, respectively), the genome assemblies of all other anemonefish species studied here are primarily based on Illumina short-read technology and are therefore highly fragmented (which results in gaps and reduced contiguity). Therefore, genome quality likely underpins why our bioinformatic analysis revealed only one *erbb3b* gene in these species, whilst PCR detected two. Nevertheless, with three copies of the *erbb3b* gene detected in *A. clarkii*, it is clear that *A. clarkii* possess a higer number than all other anemonefish species.

### Implications of an additional *erbb3b* gene in *A. clarkii*

Interestingly, alternative splice variants encoding different isoforms have been characterized for *erbb3*. Specifically, one isoform (p45-sErbB3) encoded by a 2.1 kb transcript lacks the transmembrane and cytoplasmic domains and is secreted outside the cell to modulate the activity of the membrane-bound isoform (Chen et al., 2007; Lin et al., 2008). Here, however, *erbb3b* gene1 and *erbb3b* gene2 do not seem to be the result of alternative splicing. We checked bioinformatically that reads mapped to each gene were not shared by other genes and also performed PCR with the forward primers of each short form (*erbb3b* gene1 and *erbb3b* gene2) and the reverse primer of the long *erbb3b* gene3, and did not detect any bands (Figure S8). Thus, suggesting the *erbb3b* genes in *A. clarkii* might be instead tandemly duplicated genes.

Most new proteins appear to evolve from pre-existing proteins via varying degrees of modification (Andersson et al., 2015), with gene duplication certainly being a prerequisite to acquire novel functions (Zhang, 2003). The widespread distribution of tandemly duplicated gene complexes supports a strong link between gene duplication and genetic novelty, but also gene expression (Rogers et al., 2017; Zhang, 2003). More specifically, a study (Menghi et al., 2016) showed that ErbB receptors are recurrent tandemly duplicated genes with increased expression levels. Increased receptor expression has indeed been shown to be a main mechanism of deregulation of the ErbB signaling pathways (Roskoski, 2014). There is also evidence showing that *erbb3* increases the transcription of other genes even if the cytoplasmatic pathways are not activated (i.e., truncated protein) (Bian et al., 2021). Thus, taking into account all the above, we hypothesize that the two short copies (*erbb3b* gene1 and *erbb3b* gene2) identified here could have roles in modulating the activity of the full-length protein (*erbb3b* gene3) by either (1) providing additional dimerization arms for interaction with other ErbB receptors (such as *erbb2*) and/or (2) functioning as a transcriptional co-activator of other genes necessary for the development of new melanophores.

Finally, as *erbb3b* has been shown to be essential for the development of melanophores and promoting adult pigmentation pattern metamorphosis in zebrafish (Budi et al., 2008), we also measured the expression of each *erbb3b* gene in melanic and orange *A. clarkii* juveniles.

Surprisingly, none of the three *erbb3b* genes were differentially expressed (Figure S9). Ideally, *erbb3b* expression levels would had been measured from skin tissue and not whole-body as the latter could dilute the expression signal. Unfortunately, skin tissue from the original adults used for genome sequencing was unavailable, and due to the size of the body of the juveniles, it was not possible to separate the black and orange skin. Although *erbb3* is robustly expressed in skin, it is also expressed in most epithelial tissues including the intestine and liver epithelium (Wieduwilt & Moasser, 2008). Indeed, high expression levels of all three *erbb3b* genes in *A. clarkii* were observed in the liver, gall bladder, and intestines (Figure S10), which might explain the similar expression levels between black and orange juveniles. This result does not necessarily preclude a link between the additional *erbb3b* gene *A. clarkii* posesses and the melanism polymorphism of this species. Future research should endeavor to better characterize these *erbb3b* genes and investigate their implications for melanism in *A. clarkii* and other fish.

## CONCLUSION

Here, we assembled a highly contiguous and complete chromosome-scale genome of the yellowtail clownfish *A. clarkii* using PacBio long reads and Hi□C chromatin conformation capture technologies. We annotated 25,050 protein-coding genes with 97 % completeness of conserved actinopterygian genes, the highest level among anemonefish genomes available so far. Furthermore, we identified a higher number of *erbb3b* genes in *A. clarkii* compared to other anemonefish species thus suggesting a link between this gene and the natural melanism polymorphism in *A. clarkii*. The high□quality of our genome and annotation will not only serve as a resource to better understand the genomic architecture of anemonefishes, but it will further strengthen *A. clarkii* as an emerging model organism for molecular, ecological, developmental, and environmental studies of reef fishes.

## ACKNOWLEDGEMENTS

We thank Lilian Carlu for the anemonefish drawings displayed in Figures 1 and 2. Research reported in this publication was supported by funding from OIST.

## AUTHOR CONTRIBUTIONS

TR1 (Taewoo Ryu), TR2 (Timothy Ravasi), and VL conceived the genome sequencing project. TR1 designed the overall analysis. EK and MI collected fish samples. BM and MI dissected tissues for sequencing and performed sample cross-linking. BM conducted the overall computational analysis. MH performed the phylogenetic and orthology analyses. MM and JJ collected fish for gene expression analyses. EG, CZ, SM, and JJ performed the validation assays. BM and MH wrote the manuscript with input from all authors. All authors read and approved the final manuscript.

## DATA AVAILABILITY STATEMENT

The genomic and transcriptomic sequencing reads have been deposited in the NCBI GenBank database under the BioProject ID: PRJNA813357. The chromosome-scale genome assembly has been deposited in the GenBank database under the accession number: JALBFV000000000. Genome assembly, annotation, proteome, and mitogenome for *A. clarkii* are also available in the Dryad Repository: https://datadryad.org/stash/share/odvtvEuWTbDTQ43BWODojR4gFyKGlmcB199DbikJQSc

