## Supplementary Materials for "The chromosome-scale genome assembly of the yellowtail clownfish *Amphiprion clarkii* provides insights into melanic pigmentation of anemonefish"

**SUPPLEMENTARY METHODS S1**

**Nucleic acid extraction and sequencing**

For genome sequencing, the liver of the female *A. clarkii* was extracted, snap frozen in liquid nitrogen, and stored at -80 °C. Genomic DNA was extracted using a NucleoBond HMW DNA kit (MACHEREY-NAGEL, Germany) for sequencing via two different platforms. For long-read sequencing, 10 μg of genomic DNA and a SMRTbell Express Template Prep Kit 2.0 was used to generate a 20 kb SMRTbell library according to the manufacturer’s guidelines (Pacific Biosciences, CA, USA). The library was then sequenced on the PacBio sequel II platform using a Sequencing Kit 2.0 and a SMRT cells 8 M Tray. SMRT cells using 15 h movies were used. For short-read sequencing 500 ng of genomic DNA and a Next Ultra II FS DNA Library Prep Kit (New England Biolabs, Ipswich, MA, USA) was used to prepare libraries for paired-end (151 bp) sequencing on a NovaSeq6000 platform (Illumina, San Diego, CA, USA). All sequencing was conducted at OIST sequencing center (SQC).

Liver tissue from the male *A. clarkii* was extracted, with half being snap-frozen and stored at -80 °C for Hi-C sequencing. Prior to Hi-C sequencing, DNA was crosslinked using the following protocol: (1) Small pieces of liver tissue were resuspended in 15 mL of 1 % formaldehyde solution and incubated at room temperature for 20 min with periodic mixing. (2) Glycine powder was added to the solution (to a final concentration of 125 mM) and incubated at room temperature for 15 min with periodic mixing. (3) Samples were spun down at 1,000 *g* for 1 min, the supernatant was removed, and the tissue was rinsed with Milli-Q water. (4) Tissues were ground into a fine powder using a liquid nitrogen-chilled mortar and pestle and stored at -80 °C. Cross-linked samples were shipped to Phase Genomics (Seattle, WA, USA) where a Proximo Hi-C 2.0 Kit (Phase Genomics, Seattle, WA, USA) was used to prepare the proximity ligation library and process it into an Illumina-compatible sequencing library. Paired end 150 bp sequencing of these libraries was conducted on a NovaSeq6000 platform (Illumina, San Diego, CA, USA).

Tissues from the male *A. clarkii* used for Hi-C sequencing were also used for transcriptome sequencing. The remaining half of the liver as well as the optic lobe, cerebrum, remaining brain, eye, gill, heart, gall bladder, gut, intestine, fin, muscle, and spine were extracted and stored in RNAlater (Sigma Life Science, MO, USA) at -80 °C. Total RNA was extracted from the tissue using a Direct-zol RNA Miniprep kit (Zymo Research, Irvine, CA, USA), following the manufacturers guidelines. Libraries were prepared using 400 ng of total RNA using a Next Ultra II Directional RNA Library Prep Kit (New England Biolabs, Ipswich, MA, USA). Paired end 151 bp sequencing of these libraries was conducted on a NovaSeq6000 platform (Illumina, San Diego, CA, USA). Sequencing was conducted at Okinawa Institute of Science and Technology Sequencing Centre.

Finally, total RNA from the whole body of ten *A. clarkii* juveniles (five black and five orange) was extracted to perform qPCR assays of the three erbb3b genes identified in *A. clarkii*. A TissueRuptor (Qiagen, Hilden, Germany) and Maxwell RSC simply RNA Tissue Kit (Promega, Madison, USA) were used following the manufacturer’s instructions.

**SUPPLEMENTARY DISCUSSION S1**

**Assembly and annotation of mitochondrial genome**

Whilst *A. clarkii* does not have a published genome, two mitochondrial genomes are available on NCBI (accession numbers: NC_023967.1 and AB979449.1). These mitochondrial genomes have lengths of 16,976 (Tao et al., 2016) and 16,980 bp (Thongtam Na Ayudhaya et al., 2019) respectively, and thus differ slightly in length from the mitochondrial genome here. Despite the differences in length, BLASTn v2.10.0 showed that our mitochondrial genome had sequence identities of 98.6 % (16,739 of 16,976 bp) and 97.2 % (16,501 of 16,980 bp) with these two genomes. A comparison between the other two genomes displayed a sequence identity of 98.2 % (16,673 of 16,980 bp), indicating that the mitochondrial genome published here and NCBI: NC_023967.1 are the closest match. The differences in length between these mitochondrial genomes are primarily driven by differences in the structure and length of tandem repeats in the D-loop control region. All three mitochondrial genomes contain tandem repeats, however, NC_023967.1 and AB979449.1 have five 82 bp continuity tandem repeats and one imperfect tandem repeat, whereas the mitochondrial genome assembled here has three 81 bp continuity tandem repeats and one imperfect tandem repeat. Thus, the 237 bp difference between NC_023967.1 and the genome assembled here can in part be explained by these two missing repeats. Such differences in tandem repeat structure are common in animals, with differences amongst species also being common (Omote et al., 2013; Wang et al., 2015).

**SUPPLEMENTARY FIGURES**

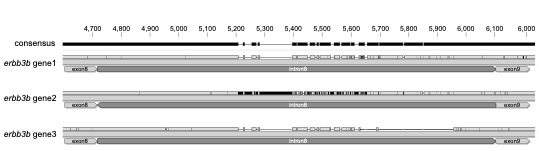

**Figure S1.** Region of intron 8 of the three *erbb3b* genes identified in *Amphiprion clarkii*. This region was chosen to design primers for polymerase chain reaction (PCR) because of the different gap lengths among genes: *erbb3b* gene1 has a gap between 5,300−5,400 whereas *erbb3b* gene2 does not have gaps but many insertions instead (represented by black lines) and *erbb3b* gene3 has two gaps between 5,300−5,400 and 5,600−6,000. Top bar shows the scale in base pairs and the consensus sequence.

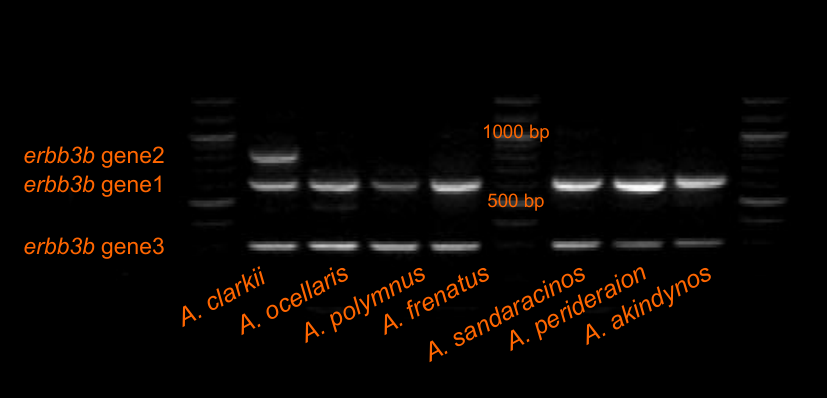

**Figure S2.** Agarose gel electrophoresis (2 %) following polymerase chain reaction (PCR) for the three *erbb3b* genes identified in *Amphiprion clarkii*. Two bands corresponding to *erbb3b* gene1 and *erbb3b* gene3 were also observed in *Amphiprion ocellaris*, *Amphiprion polymnus*, *Amphiprion frenatus*, *Amphiprion sandaracinos*, *Amphiprion perideraion*, and *Amphiprion akindynos*.

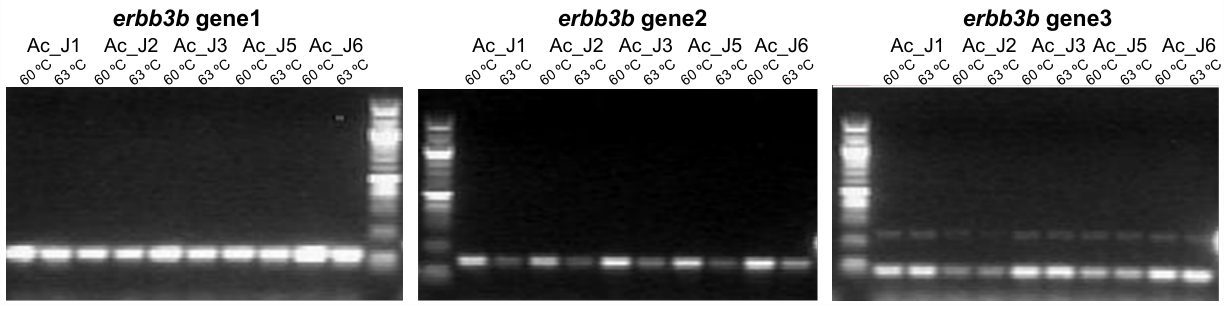

**Figure S3.** Agarose gel electrophoresis (2 %) following polymerase chain reaction (PCR) at 60 °C and 63 °C for orange (Ac_J1, Ac_J2, Ac_J3) and melanic (Ac_J5, Ac_J6) *Amphiprion clarkii* juveniles using pairs of primers targeting each of the three *erbb3b* genes.

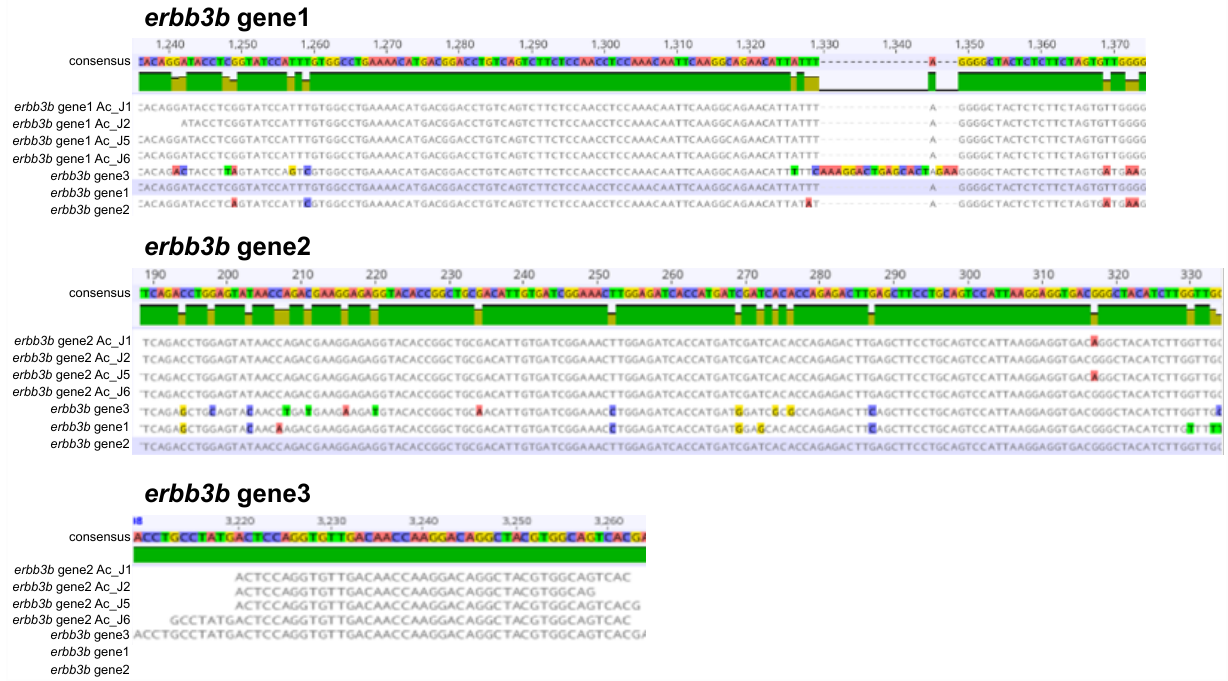

**Figure S4.** Alignment of the regions targeted by the primers for *erbb3b* gene1, *erbb3b* gene2, and *erbb3b* gene 3 after polymerase chain reaction (PCR) at 65°C in orange (Ac_J1, Ac_J2) and melanic (Ac_J5, Ac_J6) *Amphiprion clarkii* juveniles by comparison with the genomic sequences (exons only) of each gene.

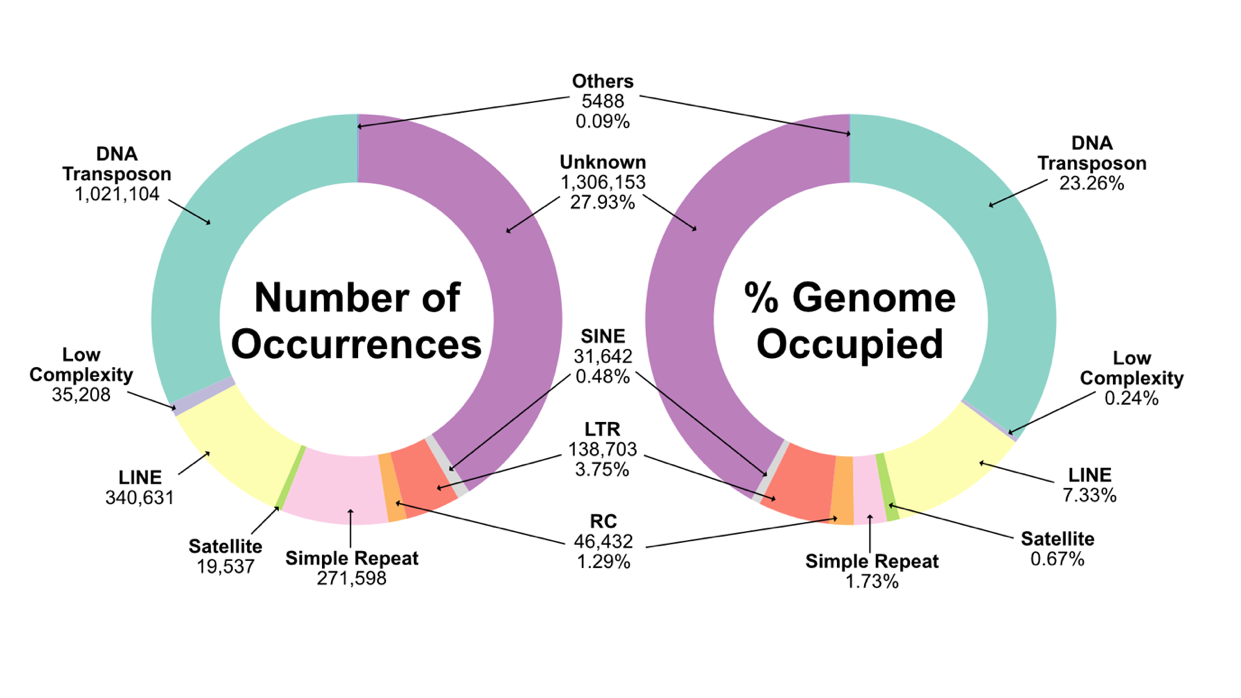

**Figure S5.** Repeat composition of the *Amphiprion clarkii* genome. Left displays the number of occurrences of each repeat type and right displays the percentage of the genome (in base pairs) occupied by each repeat type. The sum of genome percentages occupied by different types of repetitive element exceeds the total repeat content percentage of the genome as repetitive elements overlap and are nested inside each other within the genome.

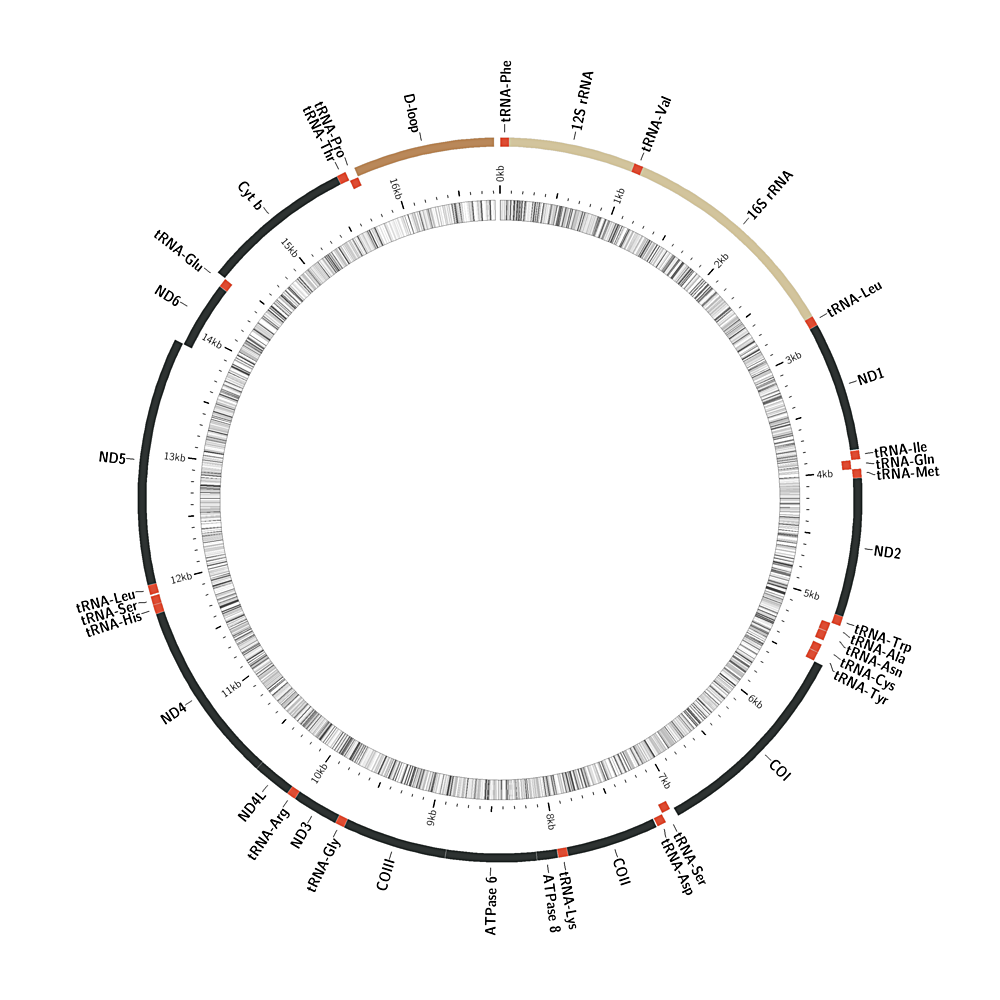
**Figure S6.** Mitochondrial genome of *Amphiprion clarkii*.

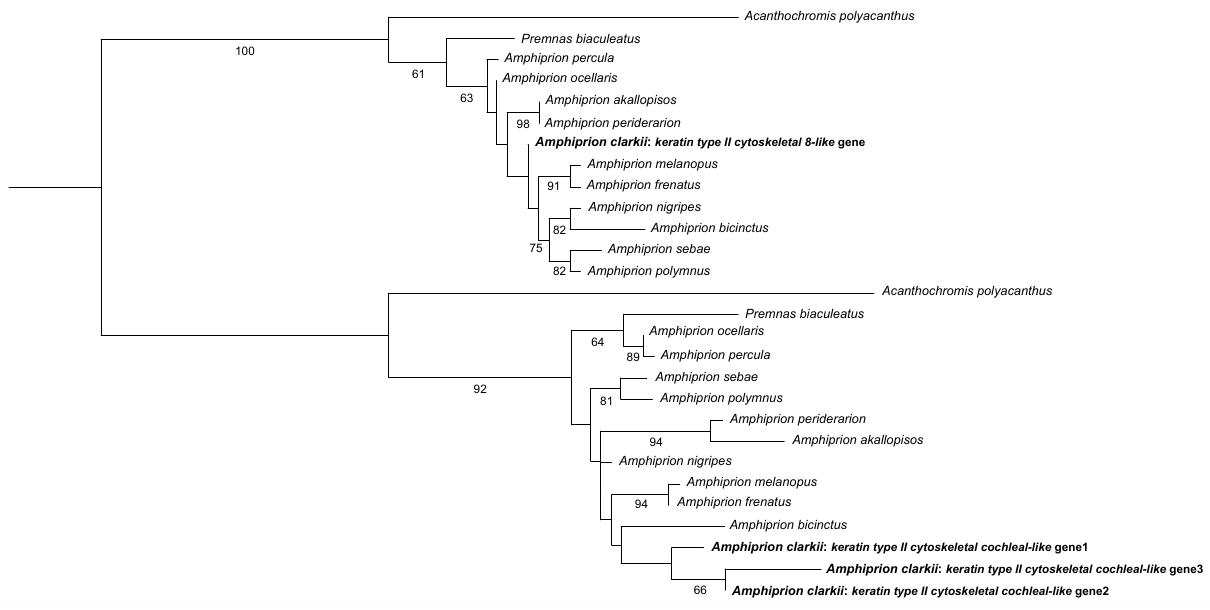
**Figure S7.** Maximum-likelihood phylogeny of protein sequences from the keratin type II cytoskeletal gene in anemonefish. Boostrap support values (%) above 50 are shown in each branching node.

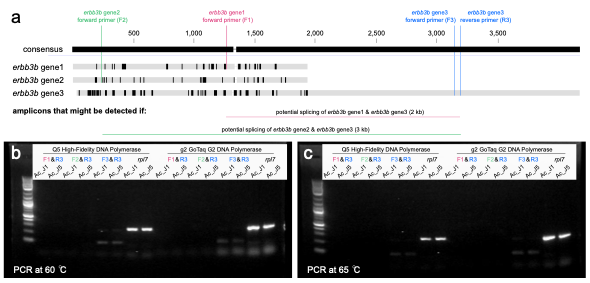

**Figure S8. (a)** Experimental setup to test if the three erbb3b genes identified in *Amphiprion clarkii* are gene duplicates or alternative splice variants. Polymerase chain reaction (PCR) assays were performed using the forward primers of each *erbb3b* gene1 and *erbb3b* gene2 (shown in red and green, respectively) with the reverse primer of *erbb3b* gene3 (shown in blue). Each gene is represented by a gray block with black lines (insertions). Top bar shows the scale in base pairs and the consensus sequence. PCRs were performed on orange (Ac_J1) and black (Ac_J5) *A. clarkii* juveniles using Q5 High-Fidelity DNA Polymerase and g2 GoTaq G2 DNA Polymerase at **(b)** 60 °C and **(c)** 65 °C.

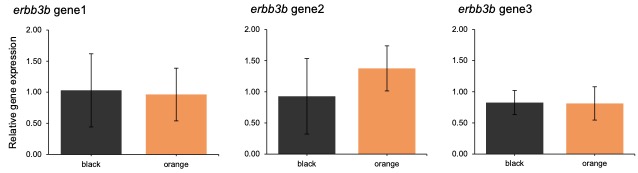

**Figure S9.** Relative expression (mean ± SD) of each *erbb3b* gene in black and orange *Amphiprion clarkii* juveniles. No differential expression was detected (unpaired two-sample Wilcoxon test: *erbb3b* gene1 W = 10, *p* = 0.69; *erbb3b* gene2 W = 18, *p* = 0.31; *erbb3b* gene3 W = 13, *p* = 1.0).

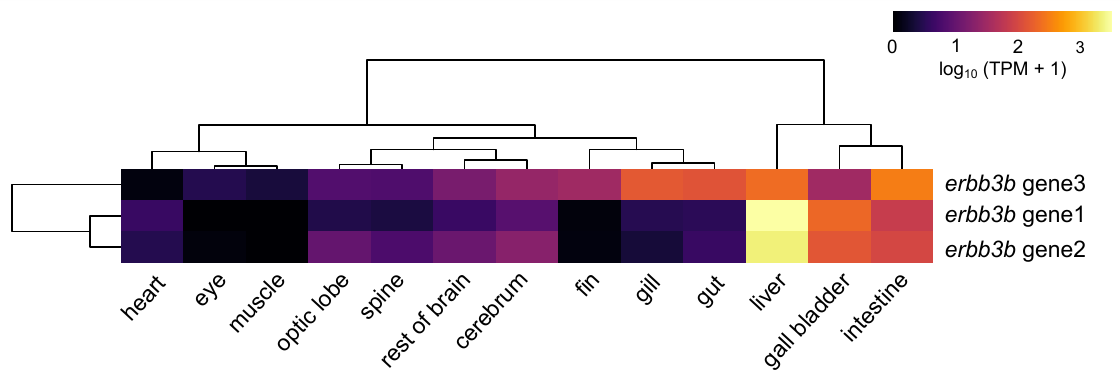

**Figure S10.** Expression levels of each *erbb3b* gene across thirteen tissues in *Amphiprion clarkii*. Color key indicates log_10_ -transformed transcript per million (TPM + 1) values.
